# A proposal for a standardized bacterial taxonomy based on genome phylogeny

**DOI:** 10.1101/256800

**Authors:** Donovan H. Parks, Maria Chuvochina, David W. Waite, Christian Rinke, Adam Skarshewski, Pierre-Alain Chaumeil, Philip Hugenholtz

## Abstract

Taxonomy is a fundamental organizing principle of biology, which ideally should be based on evolutionary relationships. Microbial taxonomy has been greatly restricted by the inability to obtain most microorganisms in pure culture and, to a lesser degree, the historical use of phenotypic properties as the basis for classification. However, we are now at the point of obtaining genome sequences broadly representative of microbial diversity by using culture-independent techniques, which provide the opportunity to develop a comprehensive genome-based taxonomy. Here we propose a standardized bacterial taxonomy based on a concatenated protein phylogeny that conservatively removes polyphyletic groups and normalizes ranks based on relative evolutionary divergence. From 94,759 bacterial genomes, 99 phyla are described including six major normalized monophyletic units from the subdivision of the Proteobacteria, and amalgamation of the Candidate Phyla Radiation into the single phylum Patescibacteria. In total, 73% of taxa had one or more changes to their existing taxonomy.

## Introduction

The rapid expansion of sequenced microbial genomes in the past few years now makes it feasible to construct a detailed taxonomy based on genome sequences (Segata et al., 2013; Hugenholtz et al., 2016; Garrity et al., 2016; Yoon et al., 2017). Microbial taxonomy, the classification of microorganisms, is essential for accurately describing microbial diversity and providing a common language for communicating scientific results (Godfray 2002). Sequence-based phylogenetic trees provide a natural framework for defining a taxonomy that takes into account evolutionary relationships and differing rates of evolution. Current microbial taxonomy is often inconsistent with evolutionary relationships as many taxa circumscribe polyphyletic groupings (McDonald et al., 2012). This is partly attributable to historical phenotype-based classification as exemplified by the clostridia where microorganisms sharing morphological similarities have been erroneously classified in the genus *Clostridium* (Yutin & Galparin, 2013; Beiko 2015). Modern microbial taxonomy is primarily guided by 16S rRNA relationships and such discrepancies are observable in 16S rRNA gene trees (McDonald et al., 2012; Yilmaz et al., 2014), but most have not been corrected due to the scale of the task and the lengthy process of formally reclassifying microorganisms (Yarza et al., 2014).

A second less obvious issue with the existing sequence-based microbial taxonomy is the uneven application of ranks across the tree. Regions that are the subject of intense study tend to be split into more taxa than other parts of the tree with equivalent phylogenetic depth, for example, the family *Enterobacteriaceae* (comprising dozens of genera) is equivalent to single genera in other parts of the tree such as *Bacillus* (Abbott & Janda, 2006). Conversely, understudied groups are often lumped together, for example, the phylum Synergistetes is currently represented by a single family (Jumas-Bilak et al., 2009) that would constitute multiple family level groupings in more intensively studied parts of the tree. A recent study by Yarza et al. (2014) proposed standardizing taxonomic ranks using 16S rRNA sequence identity thresholds and showed a high degree of discordance between these thresholds and existing taxonomy.

Current microbial taxonomies based on 16S rRNA gene relationships have a number of limitations including low phylogenetic resolution at the highest and lowest ranks (Case et al., 2007; Janda et al., 2007), missing diversity due to primer mismatches (Schulz et al., 2017), and PCR-produced chimeric sequences that can corrupt tree topologies by drawing together disparate groups (DeSantis et al., 2006). Trees inferred from the concatenation of single-copy vertically-inherited proteins provide higher resolution than those obtained from a single phylogenetic marker gene (Ciccarelli et al., 2006; Thiergart et al., 2014), and are increasingly representative of microbial diversity as culture-independent techniques are now producing thousands of metagenome-assembled genomes (MAGs) from diverse microbial communities (Brown et al., 2015; Anantharaman et al., 2016; Parks et al., 2017). Despite some caveats of their own (Thiergart et al., 2014), concatenated protein trees have been extensively used in the literature (Tonini et al., 2015; Brown et al., 2015; Hug et al., 2016), are largely congruent with 16S rRNA gene trees (Brown et al., 2001; Ciccarelli et al., 2006), and have been proposed as the best basis for a reference bacterial phylogeny (Lang et al., 2013).

Here we propose a bacterial taxonomy based on a phylogeny inferred from the concatenation of 120 ubiquitous, single-copy proteins that covers 94,759 bacterial genomes, including 13,636 (14.4%) from uncultured organisms (metagenome-assembled or single-cell genomes). Taxonomic groups within this classification describe monophyletic lineages of similar phylogenetic depths after normalizing for lineage-specific rates of evolution. This taxonomy, which we have called the GTDB taxonomy, is publicly available at the Genome Taxonomy Database website (http://gtdb.ecogenomic.org).

## Results

### Deriving the GTDB taxonomy

A dataset comprising 87,106 bacterial genomes was obtained from RefSeq/GenBank release 80 (Haft *et al.*, 2017) and augmented with 11,603 MAGs obtained from Sequence Read Archive metagenomes according to the approach of Parks et al. (2017). After removal of 2,482 of these genomes based on a completeness/contamination threshold and 1,468 based on a multiple sequence alignment threshold, the resulting 94,759 were dereplicated to remove highly similar genomes, with high-quality reference material being retained as representatives where possible (*see* *Methods*). Nearly 40% (8,559) of the dereplicated dataset of 21,943 genomes represent uncultured organisms reflecting the microbial diversity currently being revealed by culture-independent techniques (Brown et al., 2015; Anantharaman et al., 2016; Parks et al., 2017). A bacterial genome tree was inferred from the dereplicated dataset using a concatenated alignment of 120 ubiquitous, single-copy proteins (subsequently referred to as ‘bac120’; Parks et al., 2017) comprising a total of 34,744 columns after trimming 1,021 represented in <50% of taxa and 5,390 with an amino acid consensus <25%. The bac120 dataset represents ~4% of an average bacterial genome and is comparable to other bacterial domain marker sets (Dupont et al., 2012; Wu et al., 2013a).

Having inferred the concatenated protein phylogeny, we annotated the tree with group names using the National Center for Biotechnology Information (NCBI) taxonomy for the public genomes standardized to seven ranks (see *Methods*; Federhen 2012). Taxon names were overwhelmingly assigned to interior nodes with high bootstrap support (99.7% ± 2.9%) to ensure taxonomic stability. However, a few poorly supported nodes (<70%) in the bac120 tree were assigned names based on independent analyses or to preserve widely used existing classifications (**Supp. Table 1**; *see Firmicutes example below*). Since over a third of the dataset represents uncultured organisms, a substantial part of the tree was not effectively annotated using the NCBI genome taxonomy. Therefore, 16S rRNA gene sequences present in the MAGs were classified against the Greengenes 2013 (McDonald et al., 2012) and SILVA v123.1 (Yilmaz et al., 2014) taxonomies to provide additional taxonomic identifiers. Using a set of criteria to ensure accurate mapping between 16S rRNA and MAG sequences (*see* *Methods*), 74 groups lacking cultured representatives were labeled with 16S rRNA-based names, including well-recognized clades such as SAR202 (Giovannoni et al., 1996), WS6 (Dojka et al., 1998) and ACK-M1 (Zwart et al., 2003) (**Supp. Table 2**). We term all such alphanumeric names non-standard placeholders to be replaced with standard validated names in due course. Curation of the taxonomy then involved two main tasks, (i) removal of polyphyletic groups and (ii) normalization of ranks based on relative evolutionary divergence.

#### Removal of polyphyletic groups

Twenty phyla and 25 classes as defined by the NCBI taxonomy could not be reproducibly resolved as monophyletic in the bootstrapped bac120 tree (**Supp. Table 3**). Most of these were the result of a small number of misclassified genomes; however, some taxa appear to be truly polyphyletic including well-known lineages such as the Firmicutes and Proteobacteria (**Supp. Table 3**). Instability of the Firmicutes has been previously noted, primarily as a result of the Tenericutes and/or Fusobacteria moving into or out of the group (Wolf et al., 2004; Hug et al., 2016). In this prominent case, we chose to preserve the existing classification until more in-depth phylogenetic analyses are performed to resolve the issue (see rationale below). Other poorly-supported lineages such as the Proteobacteria, which is widely reported as polyphyletic using the 16S rRNA gene (Lonergan et al., 1996; McDonald et al., 2012) and protein markers (Beiko, 2011; Zhang & Sievert, 2014; Vanwonterghem et al., 2016), were conservatively divided into stable monophyletic groups. Where possible, polyphyletic taxa containing the nomenclature type retained the name, and all other groups were renamed according to the International Code of Nomenclature of Prokaryotes (Oren et al., 2015; Parker et al., 2015; *see also* *Methods*). For lower level ranks, notably genus, existing names were often retained with alphabetical suffixing to resolve polyphyly in the bac120 tree (*e.g.*, *Bacillus*_A, *Bacillus*_B, etc). Only the group containing type material (if known) kept the original unsuffixed name to indicate validity of name assignment. This serves two purposes, it preserves continuity in the literature, and avoids the necessity to propose dozens of new names for highly polyphyletic groups, although ultimately we suggest that this should be done. A total of 436 genera, 152 families and 67 orders were identified as polyphyletic in the tree highlighting important deficiencies in the current taxonomy (**Supp. Table 3**). The genus *Clostridium* was the most polyphyletic, representing 121 genera spanning 29 families, followed by *Bacillus* (81 genera across 25 families), and *Eubacterium* (30 genera across 8 families). Note, however, that these numbers were also influenced by rank normalization in some cases (*see below*).

#### Rank normalization

There is currently no accepted standardized approach for assigning taxa to higher ranks (*i.e.*, genus to phylum), although 16S rRNA sequence identity and amino acid identity (AAI) thresholds have been proposed (Hugenholtz et al., 1998; Yarza et al., 2014; Konstantinidis & Tiedje, 2005). Assignment of ranks within the NCBI taxonomy is highly variable under both these measures as they have been proposed relatively recently and have not been widely adopted (Yarza et al., 2014; Hugenholtz et al., 2016). We normalized the assignment of higher taxonomic ranks using relative evolutionary divergence (RED) calculated from the bac120 tree, which is conceptually similar to the approach used by Wu et al. (2013b). Our approach provides an operational approximation of relative time with extant taxa existing in the present (RED=1), the last common ancestor occurring at a fixed time in the past (RED=0), and internal nodes being linearly interpolated between these values according to lineage-specific rates of evolution (**Fig. 1**; *see* *Methods*). RED intervals for normalizing taxonomic ranks were defined as the median RED value for taxa at each rank ±0.1 (**Fig. 1**). This represents a compromise between strict normalization and the desire to preserve existing group names on well-supported interior nodes. Visualization of the NCBI taxonomy according to RED highlights a substantial number of over-or under-classified taxa (**Fig. 2a**). To correct these inconsistencies, taxa falling outside of their RED intervals were either reassigned to a new rank (with appropriate nomenclatural changes) or to a new node in the tree (**Fig. 2b**).

**Figure 1.**
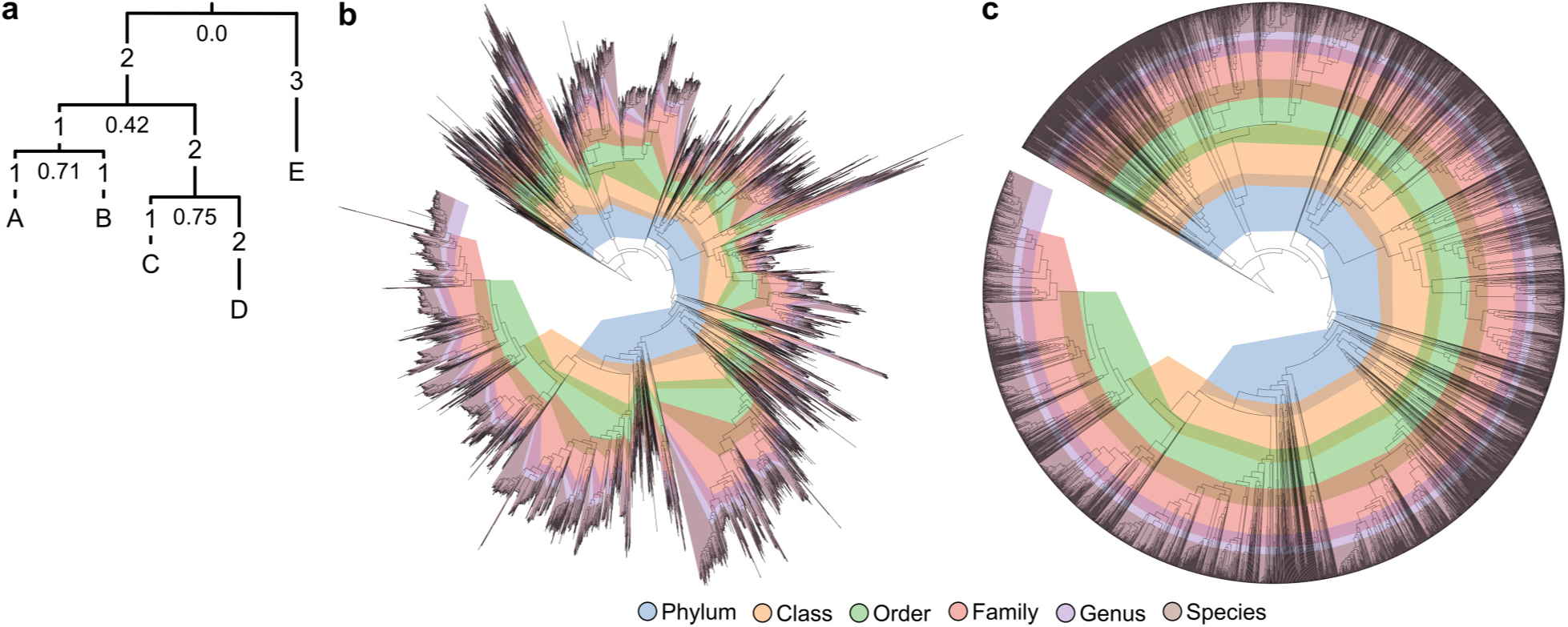
Rank normalization using relative evolutionary divergence (RED). (**a**) Example illustrating the calculation of RED. Numbers on branches indicate their length and numbers below each node indicate their RED. The root of the tree is defined to have a RED of zero and leaf nodes have a RED of one. The RED of an internal node *n* is linearly interpolated from the branch lengths comprising its lineage as defined by *p* + (*d*/*u*)×(1-*p*), where *p* is the RED of its parent, *d* is the branch length to its parent, and *u* is the average branch length from the parent node to all extant taxa descendant from *n*. For example, the parent node of leaves C and D has a RED value of 0.75 (=0.42 + (2/3.5)×(1-0.42)) as its parent has a RED of *p*=0.42, the branch length to the parent node is *d*=2, and the average branch length from the parent node to C and D is *u*=(3+4)/2=3.5. (**b**) Bacterial genome tree inferred from 120 concatenated proteins and contoured with the RED interval assigned to each taxonomic rank. Adjacent ranks overlap in some instances as this permits existing group names to be placed on well-supported interior nodes. In order to accommodate visualizing the RED intervals, the initial tree inferred across 21,943 was pruned to 8,142 genomes by retaining two genomes from each species with multiple representatives. The tree is rooted on the phylum Acetothermia for illustrative purposes. RED values used for rank normalization are averaged over multiple plausible rootings (see *Methods*). (**c**) The bacterial genome tree with branch lengths scaled by their RED values illustrating that rank normalization follows concentric rings that provide an operational approximation of relative time of divergence.

**Figure 2.**
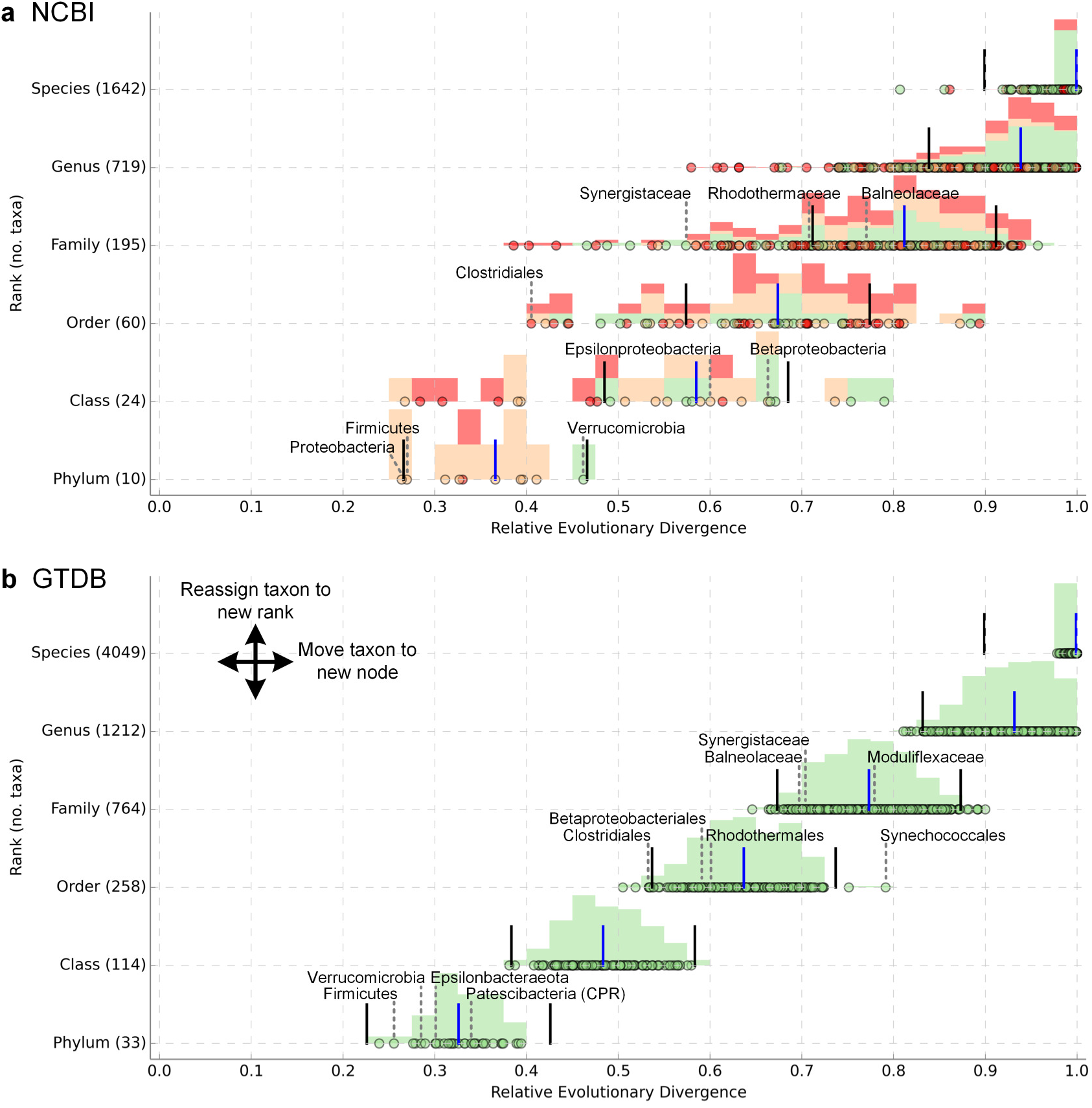
Relative evolutionary divergence of NCBI and GTDB taxa in a genome tree inferred from 120 concatenated proteins. (**a, b**) RED of taxa defined by the NCBI and GTDB taxonomies, respectively. Each point represents a taxon distributed according to their rank (Y-axis) and colored green, orange, or red to indicate the taxon is monophyletic, operationally monophyletic, or polyphyletic in the genome tree, respectively. A histogram is overlaid on the points to show the relative density of monophyletic, operationally monophyletic, and polyphyletic taxa. The median RED value of each rank is shown by a blue line and the RED interval for each rank shown by black lines. Only monophyletic or operationally monophyletic taxa were used to calculate the median RED values for each rank. The GTDB aims to resolve taxa that are over-or under-classified based on their RED value by either reassigning them to a new rank (vertical shift in plot) or moving them to a new interior node (horizontal shift in plot). For example, the family *Synergistaceae* was normalized by reclassifying the family to only encompass the genera *Synergistes*, *Cloacibacillus*, *Thermanaerovibrio* and *Aminomonas*, rather than the 12 genera circumscribed by this family in the NCBI taxonomy. Only taxa with two or more subordinate taxa are plotted as these taxa have positions in the tree indicative of their rank (*e.g.*, only 33 of the 99 phyla defined by the GTDB contain two or more classes and a phylum with a single class consisting of multiple orders is expected to have a RED value commensurate with the rank of class). The number of taxa plotted at each rank is given in parenthesis along the y-axis.

In contrast to 16S rRNA sequence identity or AAI thresholds, RED normalization accounts for the phylogenetic relationships between taxa and variable rates of evolution. For example, members of the rapidly evolving genus *Mycoplasma* (Maniloff, 2002) are sufficiently diverged to represent two phyla using a 16S rRNA gene sequence identity threshold of 75% (Yarza et al., 2014). However, vertebrate-associated *Mycoplasma* and *Ureaplasma* diverged from their arthropod-associated sister families only 400 million years ago (Maniloff, 2002), approximately consistent with the emergence of vertebrates (Kumar et al., 2017). This evolutionary event occurred much later than the primary diversification of bacterial phyla, which is estimated to have occurred between two and three billion years ago (Marin et al., 2016). The relatively recent emergence of *Mycoplasma* is more consistent with their RED-normalized ranking into a single order within the Firmicutes, than the two phyla that a 16S rRNA sequence identity of 75% would indicate.

### Validation of the GTDB taxonomy

The robustness of the approach used to generate the GTDB taxonomy was evaluated by varying marker sets, taxa, or evolutionary models. We first considered a tree inferred from a syntenic block of 16 ribosomal proteins (rp1; Brown et al., 2015; Hug et al., 2016; Parks et al., 2017) and determined the optimal position of each GTDB taxon within this tree (*see* *Methods*). On average, 94.7% of GTDB taxa at each rank were monophyletic or operationally monophyletic (defined as having an F-measure ≥ 0.95) within the rp1 tree, with the least being 92.7% at the class level and the most being 96.5% at the order level (**Fig. 3**; **Supp. Fig. 1a**; **Supp. Table 1**). Taxa that were not monophyletic within the rp1 tree were most often due to the incongruent placement of a small number of genomes, resulting in direct conflict with the GTDB taxonomy or simply unresolved as groups in the rp1 tree (*see* *Methods*). Less than 0.5% of genomes had a conflicting taxonomic assignment at any rank, and <1.5% had an unresolved taxonomic assignment at any rank with the exception of order-level assignments which were unresolved for 4.0% of genomes (**Supp. Fig. 1b; Supp. Table 4**). We also observed that taxa at the same taxonomic rank had similar RED values indicating that rank normalization was largely preserved in the rp1 tree (**Fig. 3**; **Supp. Fig. 1c**). Performing the same analysis on a 16S rRNA gene tree resulted in 78.1% (species) to 90.8% (class) of GTDB taxa being recovered as monophyletic or operationally monophyletic (**Supp. Fig. 2a**). Incongruent taxonomic assignments in the 16S rRNA tree were largely the result of unresolved taxa with <1.1% of genomes having conflicting assignments at any rank (**Supp. Fig. 2b**; **Supp. Table 5**). Taxa at the same rank had similar RED values in the 16S rRNA gene tree, though the spread of values was greater than observed on the bac120 or rp1 trees (**Fig. 3**; **Supp. Fig. 2c**).

**Figure 3.**
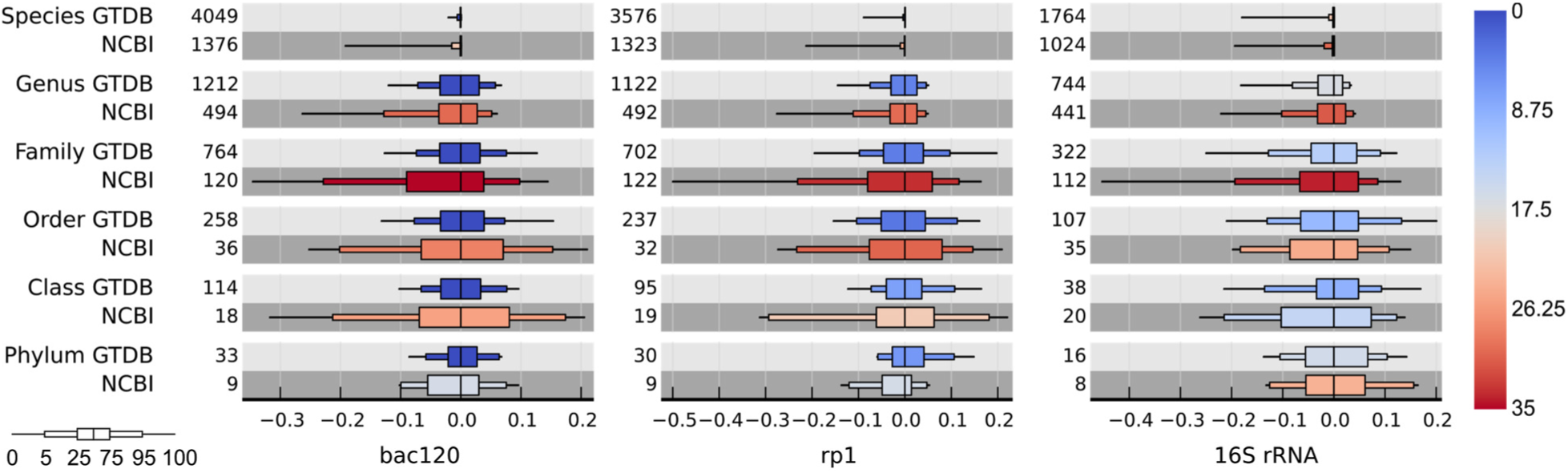
Relative evolutionary divergence (RED) and polyphyly of GTDB and NCBI taxa on trees inferred from 120 bacterial proteins (bac120), 16 ribosomal proteins (rp1), and the 16S rRNA gene. The percentage of taxa classified as polyphyletic in each tree at each rank is indicated by a color gradient from blue to red. RED distributions for taxa at each rank are shown relative to the median RED value of the rank. Results are summarized using box-and-whisker plots indicating the 0^th^/100^th^, 5^th^/95^th^, 25^th^/75^th^, and 50^th^ percentiles. Distributions were calculated over monophyletic and operationally monophyletic taxa with two or more subordinate taxa as these taxa have positions in the tree indicative of their rank. The number of taxa comprising each distribution is shown next to each box-and-whisker plot. Plots showing the RED values of individual GTDB and NCBI taxa within the bac120, rp1, and 16S rRNA trees are shown in **Fig. 2** and **Supp. Figs. 1** to **5**.

For comparison, we evaluated the congruence of the NCBI taxonomy with the topologies of the bac120, rp1 and 16S rRNA trees. All three trees had numerous discrepancies with the NCBI taxonomy both in terms of polyphyly and over-and under-classified taxa (**Figs. 2** and **3**). On average, 26.3%, 26.1% and 26.6% of NCBI taxa were classified as polyphyletic on the bac120, rp1 and 16S rRNA trees, respectively, with variable distributions across the ranks (**Supp. Figs. 3** to **5**). Curation of the GTDB on the bac120 tree ensures all taxa are monophyletic on this tree and consequently the GTDB taxa show markedly less polyphyly on the rp1 (5.3%) and 16S rRNA (14.6%) trees than NCBI taxa. In total 64.1%, 60.2% and 59.5% of RefSeq/GenBank genomes had an NCBI taxonomy congruent with the bac120, rp1 and 16S rRNA trees, while 89.8% and 76.1% of these genomes had GTDB assignments in agreement with the rp1 and 16S rRNA trees. Furthermore, the NCBI taxonomy had a poorer RED fit than the GTDB taxonomy at all ranks in these three trees (**Fig. 3**).

The stability of the GTDB taxonomy on trees inferred under marker set and taxon subsampling was also evaluated. Subsampling of the 120 bacterial marker genes was performed 100 times with 60 of the markers randomly selected for each replicate. Notably, 96.7% of GTDB taxa were classified as monophyletic in ≥90% of the replicate trees and only 10 taxa (0.11%) were classified as polyphyletic in ≥50% of replicates (**Supp. Table 1**). Given the lower phylogenetic resolution of individual genes (Gadagkar et al., 2005; Lang et al., 2013), results from individual gene trees were also remarkably robust with 86.1% of GTDB taxa being monophyletic in ≥50% of trees (**Supp. Table 1)**. Taxon resampling with one genome per genus was performed 100 times with representative genomes being randomly selected each replicate. Across the 1,430 taxa with two or more genera, 97.5% were recovered as monophyletic in ≥90% of the taxon-resampled trees and only four taxa (f__F082, f__Hyphomonadaceae, c__Bacilli_A, p__Desulfobacteraeota) were classified as polyphyletic in ≥50% of replicates (**Supp. Table 1**). The proposed taxonomy was also robust to model selection with only two GTDB taxa being operationally monophyletic (p__Bdellovibrionaeota; p__Firmicutes) and three being polyphyletic (c__Bacilli_A; o__Ryanbacterales; s__Blastomonas delafieldii) under a tree inferred with the LG protein substitution model (Le & Gascuel, 2008) instead of the WAG model (Whelan & Goldman, 2001; **Supp. Table 1**).

### Comparison of the GTDB and NCBI taxonomies

Overall, 73% of the 84,634 genomes with an NCBI taxonomy had one or more changes to their classification above the rank of species (**Fig. 4a**). These included both reclassification of taxa and filling in missing rank name information (~3% of genus to phylum names are currently undefined across the 84,634 genomes with an NCBI taxonomy). On average, 22% of names were changed per rank, the least being 7% at the phylum level, and the most being 50% at the order level (**Fig. 4a**). A total of 199 NCBI names above the rank of species were ‘retired’ from the GTDB taxonomy mostly as a result of RED normalization (**Supp. Table 6**). A detailed listing of changes overviewed in **Fig. 4a** can be found in **Supp. Table 3**. Only 18% of GTDB taxa above the rank of species were validly published and approved, a further 19% proposed but not validated (many of these lack type material), and 63% were non-standard placeholder names (**Fig. 4b**) indicating the scope of the task remaining to produce a fully standardized and validated taxonomy. This task will be greatly facilitated by recent proposals to use genome sequences as type material for as-yet-uncultured lineages, which in principle allows validation of names (Whitman 2016; Konstantinidis et al., 2017).

**Figure 4.**
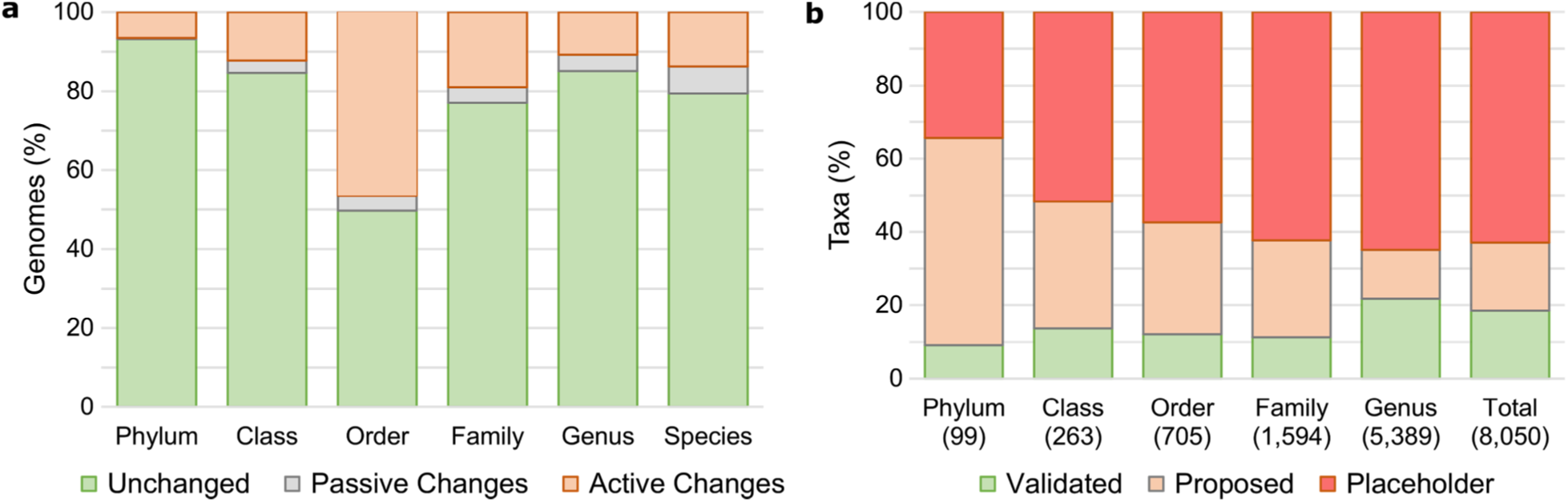
Comparison of GTDB and NCBI taxonomies and naming status of GTDB taxa. (**a**) Comparison of GTDB and NCBI taxonomic assignments across 84,634 bacterial genomes from RefSeq/GenBank release 80. For each rank, a taxon was classified as being unchanged if its name was identical in both taxonomies, passively changed if the GTDB taxonomy provided name information absent in the NCBI taxonomy, or actively changed if the name was different between the two taxonomies. (**b**) Percentage of GTDB taxa at each rank that are validly published and approved, proposed but not validated, or are non-standard placeholder names. The number of taxa at each rank is shown in parentheses.

#### Genus-and species-level classifications

Genera and species comprise 84% of the 16,924 defined taxon names in the bac120 tree. Misclassified species in the public repositories are a particular area of concern to researchers as they can introduce noise into a variety of analyses, including strain typing (Comas et al., 2009), biogeographic distributions of species (Martiny et al., 2006) and pangenome analyses (Trost et al., 2010). Moreover, classification errors can propagate over time as incorrectly labeled genomes are used as reference material to identify novel sequences. A small number of microbial genera have been rigorously examined for this problem and taxonomic corrections proposed, including *Aeromonas* (Beaz-Hidalgo et al., 2015) and *Fusobacterium* (Kook et al., 2017). We compared the results of these analyses to the GTDB taxonomy as a means of providing independent verification of our results. Based on multilocus sequence analysis and average nucleotide identity (ANI) comparisons, Beaz-Hidalgo et al. (2015) proposed that nine *Aeromonas dhakensis* genomes are incorrectly classified as *A. hydrophila*. All nine of these strains were reclassified as *A. dhakensis* in the bac120 tree, and an additional four genomes not included in the Beaz-Hidalgo study were also reclassified as *A. dhakensis* (**Supp. Table 7**). Kook et al. (2017) recently recommended the reclassification of *Fusobacterium nucleatum* subspecies *animalis*, *nucleatum*, *polymorphum*, and *vincentii* as separate species based on ANI and genome distance metrics. Rank normalization of the GTDB taxonomy using RED values largely reproduced this finding without prior knowledge of the authors’ work (**Supp. Table 7**). Reclassification of species according to the bac120 tree is also consistent with recent efforts to objectively define bacterial species based on barriers to homologous recombination estimated against the core genome of each species (Bobay & Ochman, 2017). In that study, 23 of 91 bacterial species were proposed to contain one or more members not belonging to their respective species (‘excluded taxa’). We found that almost all comparable instances of excluded taxa were due to misclassification in the NCBI taxonomy (**Supp. Table 7**). These results suggest that the bac120 tree topology and RED estimates of species-level groups based on ~4% of the genome (120 conserved markers) are consistent with alternative analytical approaches using larger fractions of the genome.

The genus *Clostridium* is widely acknowledged to be polyphyletic and efforts have been made to rectify this problem, including a global attempt to reclassify the genus using a combination of phylogenetic markers (Yutin & Galperin, 2013). These authors proposed the reclassification of 78 *Clostridium* species, and nine other species, into six novel genera (Yutin & Galperin, 2013; Galperin et al., 2016). Of these, we could confirm that *Erysipelatoclostridium* (with the exception of *C. innocuum* str. 2959), *Gottschalkia* and *Tyzzerella* (excepting *C. nexile* CAG:348) represent monophyletic genus-level groups. The remaining three genera proposed by Yutin & Galperin (2013) represent multiple genera in the GTDB taxonomy including validly published genera (**Supp. Table 8**). This is consistent with several recent analyses of individual taxa in these groups (Yarza et al., 2008; Lawson et al., 2016; Sakamoto et al., 2017). The GTDB taxonomy is also largely in agreement at the genus level with a recent global genome-based classification of the Bacteroidetes (Hahnke et al., 2016). Of the 122 genera addressed in that study, six were found to be in need of reclassification; *Chryseobacterium*, *Epilithonimonas*, *Aequorivita*, *Vitellibacter*, *Flexibacter*, and *Pedobacter*. All six were similarly identified as polyphyletic in the GTDB taxonomy and reclassified accordingly. These findings demonstrate that our methods are broadly consistent with rigorous, independent analyses of problematic genera and species.

#### Higher-level classifications

A number of notable taxonomic changes at higher ranks are proposed for well-studied groups. For example, the class *Betaproteobacteria* was reclassified as an order within the class *Gammaproteobacteria* because it is entirely circumscribed within the latter group, and is closer to the median RED value for an order than a class (**Fig. 2a**). This change is consistent with the original 16S rRNA gene topology of the Proteobacteria and subsequent trees (Ludwig & Klenk, 2001; Garrity et al., 2005; McDonald et al., 2012; Quast et al., 2013; Yilmaz et al., 2014), although such a rank change was not proposed in these studies. The *Deltaproteobacteria* and *Epsilonproteobacteria* have been removed entirely from the Proteobacteria as this phylum is not consistently recovered as a monophyletic unit, as found in many previous 16S rRNA and other marker gene analyses (*e.g.*, Rinke et al, 2013; Yarza et al., 2014; Waite et al., 2017). In the case of the *Epsilonproteobacteria*, this class was combined with the order *Desulfurellales* (*Deltaproteobacteria*) to form a new phylum, the Epsilonbacteraeota (Waite et al., 2017).

The Firmicutes also underwent extensive internal reclassification. As a clade, this phylum is typically monophyletic but poorly supported in most trees (**Supp. Table 1**), and has a RED in the phylum range, albeit to the left of the median for this rank (**Fig. 2b**). The Firmicutes have therefore been retained as a phylum-level lineage, although future revision of this status may be warranted. The taxa that comprise this phylum have been divided into 34 classes including the mycoplasmas which are currently classified as a separate phylum, the Tenericutes (Brown, 2010), and 14 classes exclusively comprised of MAGs. Incorporation of the Tenericutes within the Firmicutes is consistent with single-gene phylogenies (Wolf et al., 2004; Yarza et al., 2008; McDonald et al., 2012; Quast et al., 2013), and is further supported by recent evidence using multiple molecular markers (Lang et al., 2013; Segata et al., 2013; Hug et al., 2016; Skennerton et al., 2016). Similar to its type genus, the order *Clostridiales* has been extensively subdivided (**Fig. 5a**) largely as a consequence of an anomalous RED for this rank (**Fig. 2a**).

**Figure 5.**
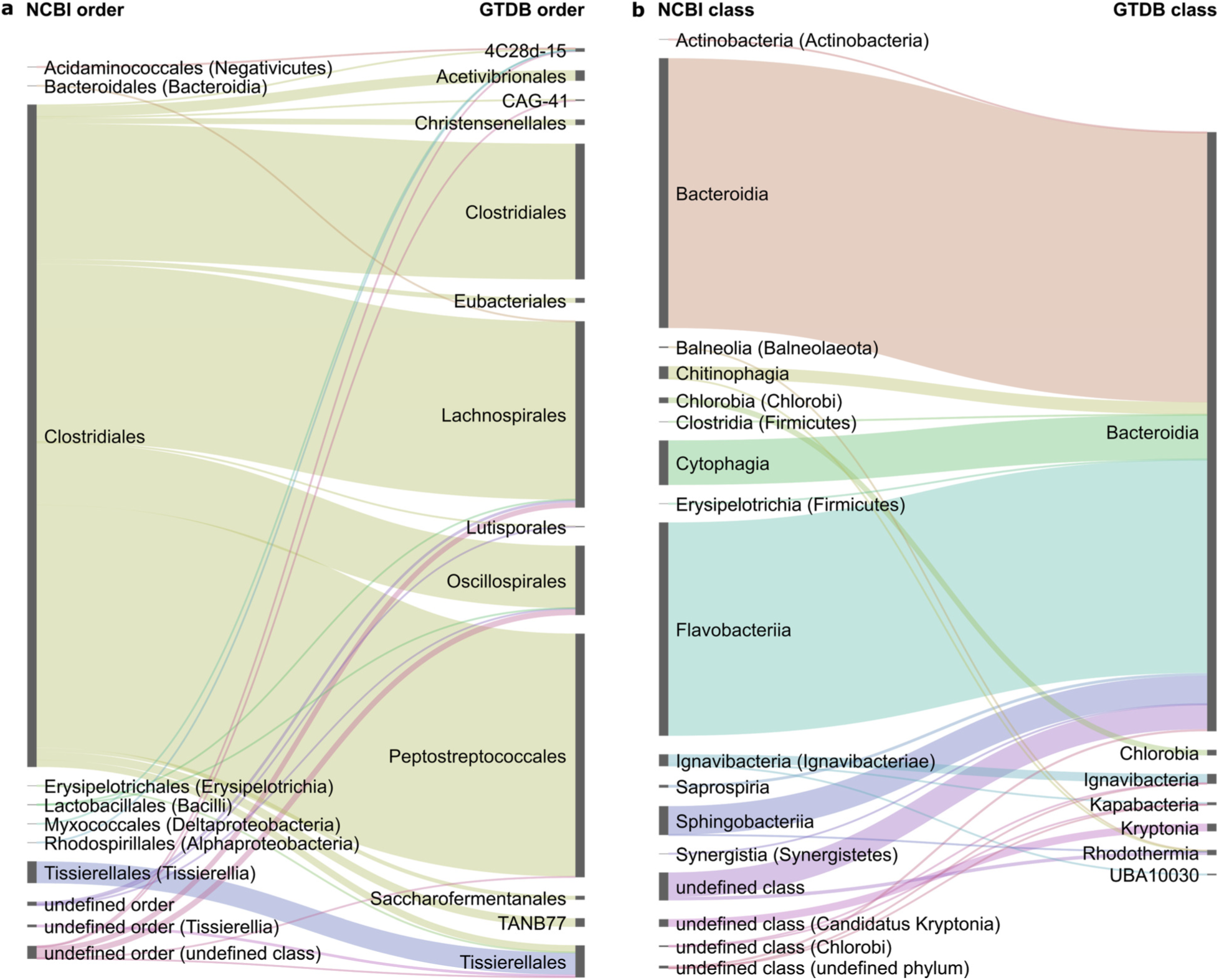
Comparisons of NCBI and GTDB classifications of genomes designated as *Clostridia* or *Bacteroidetes* in the GTDB taxonomy. (**a**) Comparison of NCBI (left) and GTDB (right) order-level classifications of the 2,368 bacterial genomes assigned to the class *Clostridia* in the GTDB taxonomy. Genomes classified in a class other than *Clostridia* by NCBI are indicated in parentheses. (**b**) Comparison of NCBI and GTDB class-level classifications of the 2,058 bacterial genomes assigned to the phylum *Bacteroidetes* in the GTDB taxonomy. Genomes classified in a phylum other than the *Bacteroidetes* by NCBI are indicated in parentheses.

Based on robust monophyly, rank normalization and naming priority in the literature, the phylum Bacteroidetes is proposed to encompass the Chlorobi (Garrity et al., 2001) and Ignavibacteriae (Podosokorskaya et al., 2013) as class-level lineages. Concomitantly, several former classes of Bacteroidetes have been amalgamated into the class *Bacteroidia* as order-level lineages including the *Chitinophagales*, *Cytophagales*, *Flavobacteriales* and *Sphingobacteriales* (**Fig. 5b**). These proposed changes are in contrast to recent reclassifications, in which Bacteroidetes is divided into three major lineages by promoting the families *Rhodothermaceae* and *Balneolaceae* to phyla (**Fig. 2a**; Hahnke et al., 2016; Munoz et al., 2016). In the GTDB taxonomy, these are retained as families within their own orders in the class *Rhodothermia* based on their RED values (**Fig. 2b**). The higher-level taxonomy of the phylum Actinobacteria is largely unchanged. The five classes *Actinobacteria*, *Acidimicrobiia*, *Coriobacteriia*, *Thermoleophilia*, and *Rubrobacteria* are retained with the sole change at the class-level being the downgrading of the *Nitriliruptoria* (Ludwig et al., 2012) to an order within the class *Actinobacteria* based on rank normalization. Changes to other major lineages are summarized in **Supp. Table 3**.

#### Calibration of uncultured microbial diversity

Having normalized the taxonomy on existing isolate-based classifications, we were able to calibrate the taxonomic ranks of uncultured lineages. Candidate phylum KSB3 was initially proposed based on comparative analysis of environmental 16S rRNA gene sequences (Tanner *et al.*, 2000; Yamada et al., 2007), and more recently two near-complete MAGs belonging to this phylum were reconstructed from a bulking sludge metagenome, for which the names *Candidatus* Moduliflexus flocculans and *Ca*. Vecturathrix granuli were proposed (Sekiguchi et al., 2015). These genomes were further classified into separate families, orders and classes within the phylum, however, by rank normalization they only represent separate genera belonging to a single family. The group still retains a phylum-level status as it is not reproducibly affiliated with other bacterial lineages (Hugenholtz et al., 1998); however, we propose that the phylum (Modulibacteria) is currently genomically represented by a single class (*Moduliflexia*), single order (*Moduliflexales*) and single family (*Moduliflexaceae*; **Fig. 2b**).

As part of a single cell genomics study, the superphylum Patescibacteria was proposed to encompass the candidate phyla Parcubacteria (OD1), Microgenomates (OP11), and Gracilibacteria (GN02) (Rinke et al., 2013). These candidate phyla were further subsumed within the Candidate Phyla Radiation (CPR) based on the addition of 797 MAGs (Brown et al., 2015). Currently there are at least 65 candidate phyla proposed to belong to the CPR (Brown et al., 2015; Anantharaman et al., 2016), with the justification of individual phyla based primarily on a 16S rRNA sequence identity threshold of 75% (Yarza et al., 2014). The CPR is consistently recovered as a monophyletic group using concatenated protein markers in this and previous studies (Brown et al., 2015; Hug et al., 2016; Parks et al., 2017). However, rank normalization suggests that the CPR should be reclassified as a single phylum for which we have reimplemented the name Patescibacteria (**Fig. 2**), with concomitant subordinate changes.

## Outlook

The GTDB taxonomy aims to provide an objective, phylogenetically consistent classification of bacterial species, and we have shown that it is largely congruent with the topology and substitution rates inferred under different gene sets and models of evolution. While we have preserved existing taxonomic classifications where possible, a substantial number of modifications were required in order to resolve polyphyletic groups and to normalize taxa at each rank based on our operational approximation of relative time of divergence. The GTDB taxonomy covers 94,759 bacterial genomes, but we expect the number of available reference genomes to expand rapidly and to quickly encompass new lineages (Anantharaman et al., 2016; Parks et al., 2017). In anticipation of this expansion, we plan to curate the taxonomy biannually in order to incorporate new genomes and proposed taxonomic groups while retaining a phylogenetically consistent classification. Subsampling of the bac120 dataset suggests that subsets of these marker genes could be used in the future to produce reliable phylogenies that better scale with the projected increase in the reference genome database (Hugenholtz et al., 2016). Some incongruencies between genome trees inferred for each biannual update are expected to impact the GTDB taxonomy as has already been observed for well-established groups such as the Firmicutes, which may require reclassification in subsequent iterations. Ideally, such regions of instability should be addressed individually with more in-depth analyses to establish the most suitable classification, as for example, was done recently with the class *Epsilonproteobacteria* (Waite et al., 2017). The GTDB taxonomy is available through the Genome Taxonomy Database website (http://gtdb.ecogenomic.org) and we are facilitating its incorporation into other public bioinformatic resources. We are also developing a standalone tool to enable researchers to classify their own genomes according to the GTDB taxonomy and its classification criteria. The methodology described here is applicable to any taxonomically annotated phylogenetic tree and we are in the process of expanding the taxonomy to Archaea and double-stranded DNA viruses. We anticipate that the availability of an up-to-date normalized genome-based classification will greatly facilitate analysis of microbial genome data and communication of scientific results.

## Methods

### Genome dataset

A dataset of 87,106 bacterial genomes was obtained from RefSeq/GenBank release 80 (Haft *et al.*, 2017) on January 17, 2017. An additional 11,603 MAGs obtained from Sequence Read Archive metagenomes (Leinonen et al., 2011) were added to this dataset to improve coverage of uncultured lineages, most of which have been reported previously (Parks et al., 2017). These genomes were dereplicated as described in Parks et al. (2017) with the exception that dereplication was based on ANI values estimated using Mash distances (Ondov et al., 2016) instead of pairwise AAI values calculated from the bac120 alignment. Specifically, a genome was assigned to a representative genome if one of the following criteria was met: i) the Mash distance was ≤0.1 (~ANI of 90%) and the genomes had the same species assignment at NCBI, ii) the Mash distance was ≤0.05 (~ANI of 95%) and the genomes had the same species assignment in the previous iteration of the GTDB taxonomy (release 78; http://gtdb.ecogenomic.org/downloads), or iii) the Mash distance was ≤0.035 (~ANI of 96.5%) and the query genomes had no species assignment at NCBI or in the previous iteration of the GTDB taxonomy. Following dereplication, genomes were excluded that had i) amino acids in <50% of the columns within the bac120 alignment and/or ii) an estimated quality <50, defined as completeness – 5 × contamination and calculated using the default lineage-specific marker gene sets of CheckM (Parks et al., 2015).

### Metadata

The NCBI taxonomy associated with the reference genomes was obtained from the NCBI Taxonomy FTP site on January 17, 2017. This taxonomy was standardized to seven ranks (domain to species) by removing non-standard ranks and identifying missing standard ranks with rank prefixes. Standard ranks were also prefixed with rank identifiers as previously described (McDonald et al., 2012). For example, the full NCBI lineage for *‘Nostoc azollae’ 0708* (GCF_000196515.1) at the time of download was “cellular organisms (no rank); Bacteria (superkingdom); Terrabacteria group (no rank); Cyanobacteria/Melainabacteria group (no rank); Cyanobacteria (phylum); Nostocales (order); Nostocaceae (family); Trichormus (genus); Trichormus azollae (species); ‘Nostoc azollae’ 0708 (strain)”, which was standardized to “d__Bacteria; p__Cyanobacteria; c__; o__Nostocales; f__Nostocaceae; g__Trichormus; s__Trichormus azollae”. Additional metadata from NCBI such as “Isolate”, “Assembly level” and “Genome representation” were also parsed from the assembly reports of all bacterial genomes (http://gtdb.ecogenomic.org/downloads) to provide information for manual tree curation. To complement the NCBI taxonomy and metadata, 16S rRNA gene sequences were identified from all NCBI genomes and MAGs using HMMER v3.1b1 (Eddy, 2011) and the most similar sequence within the Greengenes 2013 (McDonald et al., 2012) and SILVA v123.1 (Yilmaz et al., 2014) databases identified using BLASTN v2.2.30+ (Camacho et al., 2009).

### Inference and annotation of the bac120 tree

A phylogenetic tree spanning the dereplicated genomes was inferred from the concatenation of 120 ubiquitous, single-copy marker genes (bac120 marker set) identified within the Pfam v27 (Finn et al., 2014) and TIGRFAMs v15.0 (Haft et al., 2003) databases, which had previously been evaluated to be phylogenetically informative (Parks et al., 2017). Gene calling was performed with Prodigal v2.6.3 (Hyatt et al., 2010) and the 120 marker proteins identified and aligned using HMMER v3.1b1. The resulting multiple sequence alignment was trimmed by removal of columns represented by <50% of taxa and/or with an AA consensus <25%. In addition, genomes with amino acids in <50% of columns were removed before phylogenetic inference. The bac120 reference tree was inferred using FastTree v2.1.7 (Price et al., 2009) under the WAG model of protein evolution (Whelan & Goldman, 2001) with gamma-distributed rate heterogeneity (+GAMMA; Yang, 1994). Branch support was estimated by performing 100 non-parametric bootstrap replicates. Group names based on the standardized NCBI genome taxonomy were added to interior nodes of the bac120 tree using tax2tree (McDonald et al., 2012).

### Calculating relative evolutionary divergence and thresholds for taxonomic ranks

Relative evolutionary divergence (RED) values were calculated from the annotated bac120 tree using PhyloRank (v0.0.27; https://github.com/dparks1134/PhyloRank). PhyloRank performs a preorder tree traversal with the RED of the root defined to be zero and the RED of node *n* defined as *p* + (*d*/*u*)(1-*p*), where *p* is the RED of its parent, *d* is the branch length to its parent, and *u* is the average branch length from the parent node to all extant taxa descendant from *n*. As the RED of taxa is influenced by root placement and the rooting of the bacterial tree remains controversial (Williams et al., 2015), we took an operational approach and rooted trees at the midpoint of all branches leading to phyla with ≥2 classes. The RED of a taxon was then taken as the median RED over all tree rootings, excluding the tree in which the taxon was part of the outgroup. Median RED values for each taxonomic rank were determined from taxa with ≥2 immediately subordinate taxa (*e.g.*, phyla with ≥2 defined classes) and the RED rank intervals used to guide the GTDB taxonomy defined as ±0.1 from these median RED values.

### Tree-based taxonomic curation

The annotated bac120 tree was manually curated in ARB (Ludwig et al., 2004) to i) resolve polyphyletic groups, ii) correct taxa falling outside of their RED distribution and iii) add 16S rRNA-based group names to uncultured lineages. Branch lengths in the bac120 tree were replaced with their corresponding RED values to produce a ‘scaled’ tree as a visual aid in the rank normalization process (**Fig. 1c**). Polyphyletic groups were identified as part of the initial annotation of group names with numerical suffixes generated by tax2tree. Groups containing type material according to the List of Prokaryotic Names with Standing in Nomenclature (LPSN; Euzeby, 1997) kept the original unsuffixed name to indicate validity of name assignment, and other groups were renamed according to a set of nomenclatural rules (*see below*). Outlier group names (±0.1 from the median RED values) were moved into their rank distributions in one of two ways; the name was moved to another interior node in the bac120 tree, or the name was left on the original interior node but reclassified to a different rank. 16S rRNA taxonomy-based names were assigned to clades in the bac120 tree if one or more genomes spanning the clade had ≥95% identity over ≥500 bp to a reference 16S rRNA sequence with a given name. Robust interior nodes (bootstrap support >90%) were given preference for name assignments.

### Generation of final GTDB taxonomy

The GTDB taxonomy was extracted from the curated bac120 tree (Newick input format) by concatenating group names from the relevant interior nodes for each genome, and exporting them as a flat text file for validation. Validation included checks for correct number and order of ranks, presence of multiple-parents (polyphyly), orthographic and semantic errors, and consistency of order (-*ales*) and family (-*aceae*) rank suffixes. As names can only be applied to groups of ≥2 taxa in ARB, ‘singleton’ genomes often have incomplete taxonomic lineages in the exported flat text file. These were auto-completed to at least the level of genus based on the nomenclatural rules outlined below. Consistency of filled ranks between releases was tracked using additional scripts and the completed taxonomy validated once more.

### Nomenclatural rules for standard names

Nomenclatural changes of validly published names were made according to the International Code of Nomenclature of Prokaryotes (ICNP; Parker et al., 2015). In the event of the nomenclature type been excluded from or not present in the group, a new type was designated based on priority in the literature and provisional higher rank names were established with the addition of corresponding rank suffixes to the stem of the generic name. This includes the recently proposed standard suffix -*aeota* for the rank of phylum (Oren et al., 2015). Priority was established for all other taxa names, namely those without standing in nomenclature and *Candidatus* taxa, based on the earliest published taxon in the group, and ranks with missing annotations derived their name from the corresponding generic name of the earliest named taxon. The term *Candidatus* was removed from GTDB taxon names to standardize the taxonomy but is easily tracked via the NCBI Organism Name in the genome metadata (http://gtdb.ecogenomic.org). In cases where new names were not proposed to resolve polyphyly, notably for the rank of genus, alphabetical suffixes were added to the standard name (*e.g.*, *Bacillus*_A, *Bacillus*_B, etc). Species-level groups with non-standard or ambiguous names were designated as ‘genus name’ sp1, ‘genus name’ sp2, *etc*. The official naming hierarchy from lower to higher ranks was followed, with the exception of some provisional phylum names lacking named species (notably CPR phyla), which were retained after rank normalization with appropriate rank suffix changes, *e.g.* o__Levybacterales from Candidatus Levybacteria (Brown et al., 2015).

### Nomenclatural rules for non-standard placeholder names

Non-standard placeholder names were given to groups lacking standardly named representatives. Several sources were used to derive non-standard names; i) 16S rRNA environmental clone names grafted onto the bac120 tree (*see above*; **Supp. Table 2**), ii) isolate strain names, *e.g.* g__Mor1 from the genome “Acidobacteria bacterium Mor1” (GCA_001664505.1), iii) MAG names, *e.g.* g__UBA4820 from “SRA genome UBA4820” (GCA_002402325.1), and iv) genome assembly identifiers for groups exclusively comprising complex symbiont names, *e.g.* g__GCF-001602625 for “*Sodalis*-like endosymbiont of *Proechinophthirus fluctus*” (GCF_001602625.1). Non-standard names longer than 15 characters were trimmed for brevity and to minimize spelling errors, *e.g.* g__2-02-FULL-67-57 from the name “Acidobacteria bacterium RIFCSPHIGHO2_02_FULL_67_57” (GCA_001766975.1). In the rare event of identical placeholder names for two or more phylogenetically distinct groups resulting from automated name trimming, or rank filling, we appended hyphenated alphabetical suffixes (-A, -B *etc*.) to distinguish them. As with standard binomial names, species-level groups were defined as ‘genus name’ sp1, ‘genus name’ sp2 *etc*. Where necessary, non-standard names were propagated to higher ranks differentiated only by rank prefix, *e.g.* d__Bacteria; p__Acidobacteria; c__UBA4820; o__UBA4820; f__UBA4820; g__UBA4820.

### Inference of trees used to validate the GTDB taxonomy

The stability of the GTDB taxonomy was evaluated using trees inferred in a manner analogous to that described for the bac120 tree. Briefly, proteins were called with Prodigal, marker genes identified and aligned using HMMER with Pfam and TIGRfam HMMs, multiple sequence alignments trimmed based on consistency and ubiquity, genomes with poor representation in the alignment removed before phylogenetic inference, and trees inferred with FastTree under the WAG+GAMMA models unless otherwise specified.

#### Alternative Marker Sets

A ribosomal protein tree (rp1) was inferred from the concatenation of 16 ribosomal proteins (Brown et al., 2015; Parks et al., 2017) and consisted of 1,949 aligned columns after trimming 101 columns represented by <50% of taxa and 11 columns with an amino acid consensus <25%. The rp1 tree spanned 21,444 genomes after removing 1,967 genomes with amino acids in <50% of the filtered columns. Subsampled marker trees were inferred by randomly selecting 60 genes from the bac120 marker set. A total of 100 replicate trees were generated in this fashion in order to assess the impact of using different subsets of the bac120 marker set. The filtered multiple sequence alignments ranged in length from 15,010 to 20,061 amino acids and all trees spanned 21,943 genomes as no additional genome filtering was performed. Individual gene trees were also inferred for each of the genes comprising the bac120 marker set. The filtered alignments for these trees ranged in length from 43 to 1,069 amino acids and spanned 16,932 to 21,050 genomes.

#### Alternative Taxa

Trees were inferred from sets of taxa subsampled to one genome per genus in the GTDB taxonomy. Representative genomes for each genus were randomly selected from the 21,943 dereplicated genomes comprising the bac120 tree. The alignments used for the full bac120 tree were used to infer 100 taxa-subsampled trees.

#### Alternative Models

A bac120 tree under the LG (Le and Gascuel, 2008) model was inferred using FastTree v2.1.7 compiled for double precision in order to avoid numerically unstable issues. This tree was inferred using the same multiple sequence alignment as used for the bac120 tree under the WAG+GAMMA models, and spans the same set of genomes.

#### Inference of 16S rRNA gene tree

A 16S rRNA gene tree was inferred from genes >1,200 bp identified within the 21,943 dereplicated and quality controlled genomes. The 16S rRNA genes were identified using HMMER and domain-specific SSU/LSU HMM models as implemented in the ‘ssu-finder’ method of CheckM, with the longest gene being retained for genomes with multiple copies of the 16S rRNA gene. The 12,712 identified 16S rRNA genes were filtered to remove sequences potentially representing contamination using a reciprocal BLAST protocol. Genes were searched against each other using BLASTN v2.2.30+ and a gene was removed from consideration if its closest match belonged to a genome classified in a different taxonomic order as defined by the GTDB, the gene had an alignment length ≥800 bp, and the gene had a percent identity ≥82%. The percent identity criterion was based on the thresholds proposed by Yarza et al. (2014). This results in 277 sequences being removed from consideration (**Supp. Table 9**). The remaining 12,435 16S rRNA genes were aligned with ssu-align v0.1 (Nawrocki *et al.*, 2009) and trailing or leading columns represented by ≤70% of taxa trimmed which resulted in an alignment of 1,409 bp. The gene tree were inferred with FastTree v2.1.7 under the GTR (Tavaré, 1986) and GAMMA models.

### Assessing stability of the NCBI and GTDB taxonomy

The congruency of the NCBI and GTDB taxonomies on different trees was assessed by placing each taxon on the node with the highest resulting F-measure. The F-measure is the harmonic mean of precision and recall, and has been previously proposed for decorating trees with a donor taxonomy (McDonald et al., 2012). Taxonomic stability was assessed as both a percentage of taxonomic groups identified as being monophyletic, operationally monophyletic, or polyphyletic within a tree and the percentage of genomes in the tree with identical, unresolved, or conflicting taxonomic assignments relative to the NCBI or GTDB taxonomies. Since a few incongruent genomes (resulting from phylogenetic incongruence, phylogenetic instability, chimeric artifacts, or erroneous NCBI taxonomic assignments) are sufficient to cause a large number of polyphyletic taxa, we classified taxa with an F-measure ≥0.95 as operationally monophyletic. Results were restricted to taxonomic groups containing two or more genomes as taxa represented by a single genome are guaranteed to be monophyletic in a tree. Genomes in a tree with incongruent taxonomic assignments were classified as either i) *conflicting* if the genome was assigned to a different taxon, or ii) *unresolved* if the taxon had no taxonomic label at a rank or multiple taxonomic labels at a rank, one of which was the expected label (*e.g.*, a polyphyletic lineage spanning two or more genera, one of which is the expected genus for the genome).

### Data availability

Data files for the GTDB taxonomy are available at http://gtdb.ecogenomic.org and include: i) flat file with the GTDB taxonomy defined for 94,759 genomes; ii) bootstrapped bac120 tree in Newick format spanning the 21,943 dereplicated genomes and annotated with the GTDB taxonomy; iii) FASTA files for each marker gene and the trimmed concatenated alignment; iv) metadata for all genomes including NCBI, Silva, and Greengenes taxonomies, completeness and contamination estimates, assembly statistics (*e.g.*, N50), and genomic properties (*e.g.*, GC-content, genomes size); v) FASTA file of 16S rRNA gene sequences identified within the 21,943 dereplicated genomes; and vi) an ARB database containing the bac120 tree. The 3,087 MAGs introduced in this study are available under BioProject PRJNA417962 and on the GTDB website.

### Code availability

RED values were calculated with PhyloRank (https://github.com/dparks1134/PhyloRank) which is freely available under the GNU General Public License v3.0.

## Acknowledgements

We thank Pelin Yilmaz for helpful discussions on the proposed genome-based taxonomy, QFAB Bioinformatics for providing computational resources, and members of ACE for beta-testing GTDB. The project is primarily supported by an Australian Research Council Laureate Fellowship (FL150100038) awarded to PH.

## Supplementary Material

Methods and Materials

Supp. Tables 1 to 9

Supp. Figures 1 to 5

